# A geminivirus AC5 protein interacts with plant hormonal signalling and impacts plant defence

**DOI:** 10.1101/2023.07.25.550498

**Authors:** Rohit Kumar, Indranil Dasgupta

## Abstract

Geminiviruses are a large group of plant viruses responsible for yield loss in various crops, mainly in the tropical and sub-tropical regions. Geminiviruses encode six to nine multifunctional proteins, which interact with plant components to cause pathogenesis. One of the least studied geminiviral proteins is AC5. This study presents the first evidence of an AC5 protein interacting with a component of the abscisic acid signalling pathway, resulting in a depressed state. We show that the AC5 protein, encoded by Sri Lankan cassava mosaic virus is important for symptom development and virus accumulation in the experimental host *Nicotiana benthamiana*. The above interaction perturbs the abscisic acid signalling pathways, leading to compromised expression of defense-related genes and insensitivity to abscisic acid in transgenic Arabidopsis plants. This suggests a novel role of AC5 to facilitate virus propagation. Furthermore, we show that transiently suppressing the expression in *N. benthamiana PP2C* with which AC5 interacts, results in a reduction in viral titers possibly due to augmented ABA signaling and its defense-related roles. This research provides valuable insights into how geminiviral proteins manipulate ABA-mediated defence pathways, paving the way for further investigation into the underlying mechanisms and potential applications in plant protection against viral infections.

## Introduction

Hormones are instrumental in regulating plant responses to biotic and abiotic stresses, orchestrating an intricate network that modulates the plant’s response to diverse stimuli. Among these, four key hormones act as primary regulators of plant defence against pathogenic agents: salicylic acid (SA), jasmonic acid (JA), ethylene (Et), and Abscisic acid (ABA). This isoprenoid hormone has a synthesis which commences in the chloroplast with carotene (C40) and concludes in the cytosol (Xiong and Zhu, 2003, Finkelstein, 2013). ABA can be detected in several organelles, such as the chloroplast, the plasma membrane (associated with G protein-coupled receptors) and the cytosol (Guo *et al*., 2011; Dittrich *et al*., 2019). The downstream components of ABA receptors play important roles which determine the responses of plants to stress/stimulation. The physiological effects of ABA are carried out by its three chief signalling components: Pyrabactin Resistance (PYR)/Pyrabactin Resistance 1-Like (PYL)/ Regularly Component of ABA Receptors (RCAR) ABA receptors, phosphatases (PP2Cs) and SNF1-related protein kinases (SnRKs). The PP2Cs are known to fulfil a key role in the negative regulation of ABA signalling. PYR/PYL/RCAR (PYLs) proteins interact with PP2Cs and bind to ABA, releasing the inhibition of ABA-activated protein kinases (OST1/SnRK2.6/SnRK2.2/2.3, Fujii and Zhu, 2009; Fujii *et al*., 2009). This, in turn, leads to the phosphorylation and activation of downstream targets such as ABF transcriptional factors, activating or inhibiting the expression of nuclear ABA-responsive genes. Moreover, GHR1 has been demonstrated to interact with PYLs (Hua *et al*., 2012), and SnRK1 is also implicated in this signalling pathway (Rodrigues *et al*., 2013). Additionally, PP2Cs have been proposed to act as co-receptors in ABA signalling, as their binding to PYLs enhances their ABA-binding affinity (Melcher *et al*., 2009; Yin *et al*., 2009).

The involvement of ABA in interactions between plants and fungi or bacteria is diverse, and whether it acts as a stimulant for the pathogen or as an immunizing agent is dependent on the timing of its activation and the type of pathosystem (Ton *et al*., 2009). There exists a paucity of research on the precise involvement of ABA in plant-virus interactions, although extant studies support the notion that it confers enhanced resistance to various RNA viruses in dicotyledonous plants (Alazem, Kim and Lin, 2019).

In compatible plant-virus interactions, it has been observed that some dicotyledonous plants become more resilient to drought following infection with certain RNA viruses (Alazem and Lin, 2015). For instance, when various plants were infected with pathogens such as tobacco mosaic virus (TMV), cucumber mosaic virus (CMV) and brome mosaic virus, their responses related to drought stress were delayed in comparison to those of healthy plants (Xu *et al*., 2008; Alazem, Lin and Lin, 2014). Similar findings were reported when resistant soybean plants were infected with soybean mosaic virus or *N. benthamiana* plants with CMV or bamboo mosaic virus (BaMV) (Seo, Lee and Kim, 2009). This increased tolerance is usually attributed to the higher levels of ABA and/or osmoprotectants present (Finkelstein, 2013). Mutations in the ABA pathway have been found to make plants more prone to potato virus X (PVX), BaMV and the crucifer infecting TMV strain, playing a unique role in allowing the enhanced accumulation of BaMV and CMV (Chen *et al*., 2013; Alazem, Lin and Lin, 2014; Alazem and Lin, 2017). These discoveries highlight that, in these cases, ABA induction is in fact a defence response.

Studies on the role of ABA in defending against viruses have been documented, with evidence of its involvement in the regulation of plasmodesmata (PD) and the antiviral RNA silencing pathway (Alazem and Lin, 2017). In plants harbouring a specific antiviral resistance (R)-gene, a series of defence mechanisms are initiated in sequence and/or in a hierarchical fashion to remove the infecting virus (Baebler *et al*., 2014; Alazem *et al*., 2018). However, even in the absence of (R)-genes, these mechanisms can still be instigated in response to viral infection, though typically to a weaker extent and without generating robust resistance (Baebler *et al*., 2011, Baebler *et al*., 2014). By forming a callose barrier at PD, viral spread is restricted to the point of infection, thereby trapping the virus and allowing for other responses, such as RNA silencing and autophagy, to act in removing viral RNAs and proteins (Seo et al., 2009; Alazem *et al*., 2019; Wu *et al*., 2019; Reagan and Burch-Smith, 2020). It is hence apparent that ABA takes part in multiple resistance steps, and although it counters the conventional antiviral hormone SA, both ABA and SA appear to be effective in controlling the antiviral RNA silencing pathway (Alazem *et al*., 2019).

Geminiviruses belong to a large family of single-stranded DNA plant viruses, having the propensity to infect a multifarious range of plant species and causing extensive agricultural impairment (Rojas *et al*., 2015). In line with present-day taxonomic paradigms, the family *Geminiviridae* constitutes one of the largest of plant viruses with fourteen genera namely *Becurtovirus, Begomovirus, Capulavirus, Citlodavirus Curtovirus, Eragrovirus, Grablovirus, Maldovirus, Mastrevirus, Mulcrilevirus, Opunvirus, Topilevirus, Topocuvirus,* and *Turncurtovirus* and more than 520 species, based on genome structure and insect vectors (Walker *et al*., 2021; Li *et al*., 2022). The largest genus, *Begomovirus*, contains approximately 450 species having monopartite or bipartite genomes, harbouring six or nine genes. In geminiviruses, the genetic component comprises one (monopartite) or two (bipartite) DNA components that encode 5–7 proteins namely, C1-C5, V1 and V2 for monopartite and AC1-AC5, AV1, AV2, BC1 and BV1 for bipartite, which play important roles in functions essential for viral replication, movement, transmission, and pathogenesis. These viral proteins display multifunctionality, and some have undergone adaptations to perform diverse roles for different viruses, even among closely related species (Devendran *et al*., 2022).

For AC5, limited information is available on its role in the infection process. For instance, studies exploring the AC5 protein of watermelon chlorotic stunt virus (WmCSV) have indicated that it is not essential for WmCSV infectivity in plants (Kheyr-Pour *et al*., 2000). Conversely, the AC5 protein of mungbean yellow mosaic India virus (MYMIV) has been shown to perform pivotal functions in MYMIV infection in plants and exhibits RNA silencing suppression activity (Li *et al*., 2015). Correspondingly, null mutants of tomato chlorotic mottle virus (ToCMoV) AC5 have exhibited infectivity, providing further corroboration that AC5 is not indispensable for infectivity. On the other hand, functional analyses of the AC5/C5 protein in tomato yellow leaf curl virus and ageratum leaf curl Sichuan virus (ALCScV) have indicated that it is an indispensable virulence factor and is implicated in the suppression of the post-transcriptional gene silencing defence pathway (Li *et al*., 2021; Zhao *et al*., 2022).

Recent study by Zhao et al. (2023), investigated the movement of monopartite geminiviruses in plants. The TYLCV-C5 protein was shown to localize at PD and travel along microfilaments. The V2 protein is found to anchor at PD and C5 can target V2 to plasmodesmata, indicating a conserved mechanism for viral intercellular movement. C5 acts as a bridge, facilitating the transport of the V2-CP-viral DNA complex from the nucleus to plasmodesmata. This study provided insights into the molecular mechanisms underlying viral movement and highlights the cooperative role of C5 and V2 in promoting viral spread in plants. However, despite these investigative attempts, our comprehension of the range of functions exhibited by the AC5/C5 protein remains constrained.

The root crop cassava (*Manihot esculenta* Crantz) is an important source of dietary and industrial starch in several countries in the African and Asian continents (Amelework and Bairu, 2022). Sri Lankan cassava mosaic virus (SLCMV) is the predominant bipartite begomovirus found in cassava mosaic disease-affected cassava in India (Patil *et al*., 2005). Cloned DNA of SLCMV can be easily inoculated using agrobacterium onto the laboratory host *Nicotiana benthamiana* giving rise to symptoms such as stunting, leaf curling/deformation and vein swelling (Mittal *et al*., 2008). The viral DNA and transcripts accumulate in newly emerged leaves within one or two weeks, which can be used as a convenient marker for the degree of infection (Mittal *et al*., 2008). As part of our effort to understand the gene functions of SLCMV, we have used the infectious clone mentioned above to explore the functions of AC5 protein. We have used mutagenesis to show that SLCMV AC5 plays a crucial role in pathogenesis in *N*. *benthamiana*. We show that SLCMV AC5 over-expressing plants display ABA insensitivity. We have also used multiple approaches to show that ABA-INSENSITIVE1 (ABI1), a crucial PP2C acts as the interacting partner of SLCMV AC5 and silencing ABI1 in *N*. *benthamiana* compromises the pathogenicity of SLCMV. Our work reveals a new role of ABI1-AC5 interaction in modulating the virus resistance in plants.

## Material and methods

### Plant Growth conditions

Germination of wild type *N. benthamiana* seeds was carried out on a sowing medium comprised of a three-to-one proportion of coco-peat and vermiculite. After three weeks, the individual plantlets were transplanted into containers and cultivated in a growth chamber maintained at a temperature of 26 ± 2 °C with a sixteen-hour photophase and eight-hour scotophase. Plants of *Arabidopsis thaliana* ecotype Columbia (Col-0) and transgenic plants expressing 35S::AC5 were grown in pots on a perlite/soil mixture at 23°C under long-day conditions with 16-h light.

### Sequence and Bioinformatics Analysis

AC5 protein sequences were retrieved from NCBI database which was further analysed. Sequence Demarcation Tool software was used for pairwise identity scores using a color-coded matrix. For cluster analysis, CLUSTAL W alignment program of the MEGA 7 software was used. The generated pairwise similarity matrix was used to group protein sequences by the unweighted pair group method arithmetic average (UPGMA). The analysis involved 150 protein sequences.

### Construction of SLCMV_AC5m

We designed primers for mutagenesis: Mutation of 1 nucleotide to change four ATG codon to GTG codon sequence in AC5 gene in SLCMV infectious clone (Mittal *et al*., 2008). Table S1 includes four sets of representative primer sequences we used for site-directed mutagenesis.

### Agroinfiltration

For inoculation of plants, *Agrobacterium tumefaciens* cultures carrying infectious clones of SLCMV were infiltrated into *N. benthamiana* leaves at the four- to five-leaf stage. SLCMV AC5 was cloned in the PVX vector using primers containing sites of restriction enzymes *Cla*I and *Sal*I (Supplementary table S2). For recombinant PVX vector expressing wild-type form of the AC5 gene (PVX-AC5), each Agrobacterium culture was adjusted to an optical density (OD_600_) of 0.5 before infiltration into *N. benthamiana*.

### RNA quantification

Total RNA from inoculated *N*. *benthamiana* was extracted using RNeasy micro kit (QIAGEN). After DNase I treatment, the first-strand complementary DNA (cDNA) was synthesized from 1 μg of RNA using oligo (dT) primers with Reverse Transcriptase (Agilent). Quantitative reverse transcription polymerase chain reaction (qRT-PCR) was performed with the SYBR Green Supermix (Bio-Rad, Hercules, CA, USA) by a Bio-Rad CFX96™ Real-Time PCR System according to the manufacturer’s manual. Briefly, 100 ng cDNA was used as template for amplification and the relative expression levels of target genes were quantified with the calculation with transcript levels of ubiquitin and actin as an internal control. All experiments were performed three times independently.

For the quantification of viral titers and the estimation of SLCMV CP transcript, the newly emerged symptomatic leaves from inoculated plants were collected at 15 dpi, and RT-qPCR was carried out. RT-qPCR for SLCMV quantification was done using CP primers (Accession no. AJ579307) to estimate CP transcript.

For quantification of defense related gene and AGO expression in transgenic Arabidopsis expressing 35S::AC5, we deployed RT-qPCR using primers mentioned in Table S2 (Supplemental data).

### Subcellular localization

For subcellular localization, coding sequences of AC5 was introduced into pSITE4CA (Chakrabarty *et al*., 2007) by gateway cloning technology. Transient expression of pSITE4CA::AC5, PIP2::CFP (positive control) and pSITE4CA (negative control) was carried out by agroinfiltration. The infiltrated plants were cultured at 24 °C under a 16-h light/8-h darkness photoperiod. Plasmolysis of *N. benthamiana* leaf tissue was performed by immersing pieces of leaves in 20% NaCl for 10 min, described earlier (Ricachenevsky *et al*., 2018). It was followed by confocal microscopic analysis. Fluorescence in leaves of *N. benthamiana* was monitored at 3 d post-agroinfiltration, and then imaged directly using a confocal laser scanning microscope (Leica). The excitation wavelengths used were 488 nm for GFP and 587 nm for CFP.

### Western blotting and Co-immunoprecipitation assay

Different combinations of agrobacterium harbouring vectors containing AC5-myc and ABI1-FLAG which were cloned using gateway technology were infiltrated into *N. benthamiana* leaves. After 3 days, agroinfiltrated leaves were harvested and used for Co-IP assays. For Co-IP experiments, we employed protocol according to a previous report (Win *et al*., 2011).

### Bimolecular fluorescence complementation assay

For the BiFC assay, SLCMV AC5 and arabidopsis ABI1 genes were cloned into the pSYPNE and pSYPCE vectors with a YFP conserved domain to generate N-terminal or C-terminal fusion proteins. The plasmids were introduced into Agrobacterium tumefaciens GV3101 and combinations pSPYNE-ABI1/pSPYCE-AC5 and pSPYCE-ABI1, pSPYNE-AC5 were used for experiment and combinations of pSPYNE-ABI1/pSPYCE, pSPYCE-AC5/pSPYNE and pSPYNE/pSPYCE was used as negative control. After infiltration into *N. benthamiana*. Infected tissues were imaged after 48 h using a Leica SP8 confocal microscope.

### Virus-Induced Gene Silencing in *N. benthamiana*

TRV based VIGS system was used for gene silencing in *N. benthamiana*. To study antiviral role of ABI1 ortholog in *N. benthamiana*, ABI1 was aligned against *N. benthamiana* genome using solgenomics BLAST (solgenomics.net/tools/blast/?db_id=330) to find putative conserved PP2C genes. Six significantly similar sequences with more than 76% identity were selected and analysed further. Conserved 200 nt sequence among those six sequences was amplified using forward primer and reverse primer (with restriction sites *Xho*I and *Xba*I respesctively) and cloned into TRV2 vector in an antisense orientation to form TRV::Anti-PP2C. This construct was subsequently transformed into agrobacterium strain GV3101. Thereafter, the mixed agrobacterium cultures of pTRV1 and TRV:Anti-PP2C, were infiltrated into the leaves of 3-leaf-stage *N. benthamiana* plants along with infectious clone of SLCMV (Mittal, Borah and Dasgupta, 2008). The combination of TRV:00 (pTRV1 and pTRV2) was used as a negative control for the VIGS experiments whereas TRV::PDS was used as positive control for VIGS assay.

### Arabidopsis transformation and measurement of seed germination and salt tolerance

35S::AC5 construct was introduced into Agrobacterium tumefaciens strain GV3101 and used in the transformation of Arabidopsis Col-0 plants. Transgenic Arabidopsis lines were generated by Agrobacterium mediated transformation using the floral dip method (Clough and Bent, 1998). T1 transgenic seeds were selected on half-strength Murashige and Skoog medium (1/2 MS) medium containing 50 mg/L hygromycin, and T3 homozygous progeny were used for further study. At least two independent lines of each transgenic material were used in this study.

For Arabidopsis seeds plating, seeds were surface sterilized by using liquid methods. The seeds first treated with 70% ethanol for 2 min and the supernatant was discarded; then, with 10% sodium hypochlorite containing 0.3% Tween-20 for 5 min, and the supernatant was discarded, followed by rinses with sterile water four times, centrifugation for 2 min at 4,000 g each time, and the supernatant was discarded. Finally, the seeds were suspended in 0.15% sterile agarose and kept at 4°C for 2 days. Sterilized seeds were plated on plates and grown in a growth chamber at 22°C under 16 h 60–80 μE m–2 s–1 continuous light and 8 h darkness. The germination and salt tolerance phenotype of Col-0 and 35S::AC5 transgenic Arabidopsis seeds were performed by growing them on 1/2 MS supplemented with specific concentration of ABA or NaCl. Germination was determined based on the appearance of the embryonic axis. Salt tolerance phenotype was based on the observation of green cotyledons in a seedling. Three independent experiments were performed, and similar results were observed. ABA was dissolved in dimethyl sulfoxide.

### Yeast two hybrid assay

Plasmids were transformed into Y2HGold competent yeast cells using the LiAc-PEG method (Gietz and Schiestl, 2007). The transformants were selected on SD-Trp plates and the interactions were tested on SD-Leu-Trp-His plates. The primers are listed in Table S2.

## Results

### Sequence analysis of AC5 protein among different begomoviruses

To examine the sequence variation of AC5 proteins from different begomoviruses, 150 randomly selected AC5/C5 amino acid sequences were retrieved from NCBI data base and analysed for protein sequence and length variation. The alignment of amino acid sequences of AC5 indicated that it exhibits variablity, both at the number of amino acids as well as their sequence (Fig 1a and b). Pairwise analysis of the AC5/C5 amino acid (AA) sequences (n=150) with SLCMV AC5 showed predicted identity values ranging between 3% to 85% with lowest values observed with tomato golden vein virus AC5 (accession QMV80596) and highest with AC5 protein of African cassava mosaic virus (accession ACF42182) and Indian cassava mosaic virus (accession Q08591) proteins. The predicted size of the AC5 proteins ranged from 56 AA in tomato leaf curl Thailand virus to 257 AA in Indian mungbean yellow mosaic virus (Fig 1b). Phylogenetic analysis was performed on 130 C5/AC5 sequences and Sequence Identity dendrogram of AC5/C5 suggested that SLCMV AC5 and those from African cassava mosaic virus-Ghana, tomato yellow leaf curl Thailand virus (TYLCTHV), vernonia yellow vein virus, watermelon chlorotic virus, African cassava mosaic virus, East African cassava mosaic Cameroon virus, pouzolzia golden mosaic virus and East African cassava mosaic virus shared a single clade, indicating common evolutionary origin (Fig 1c).

**Figure 1.**
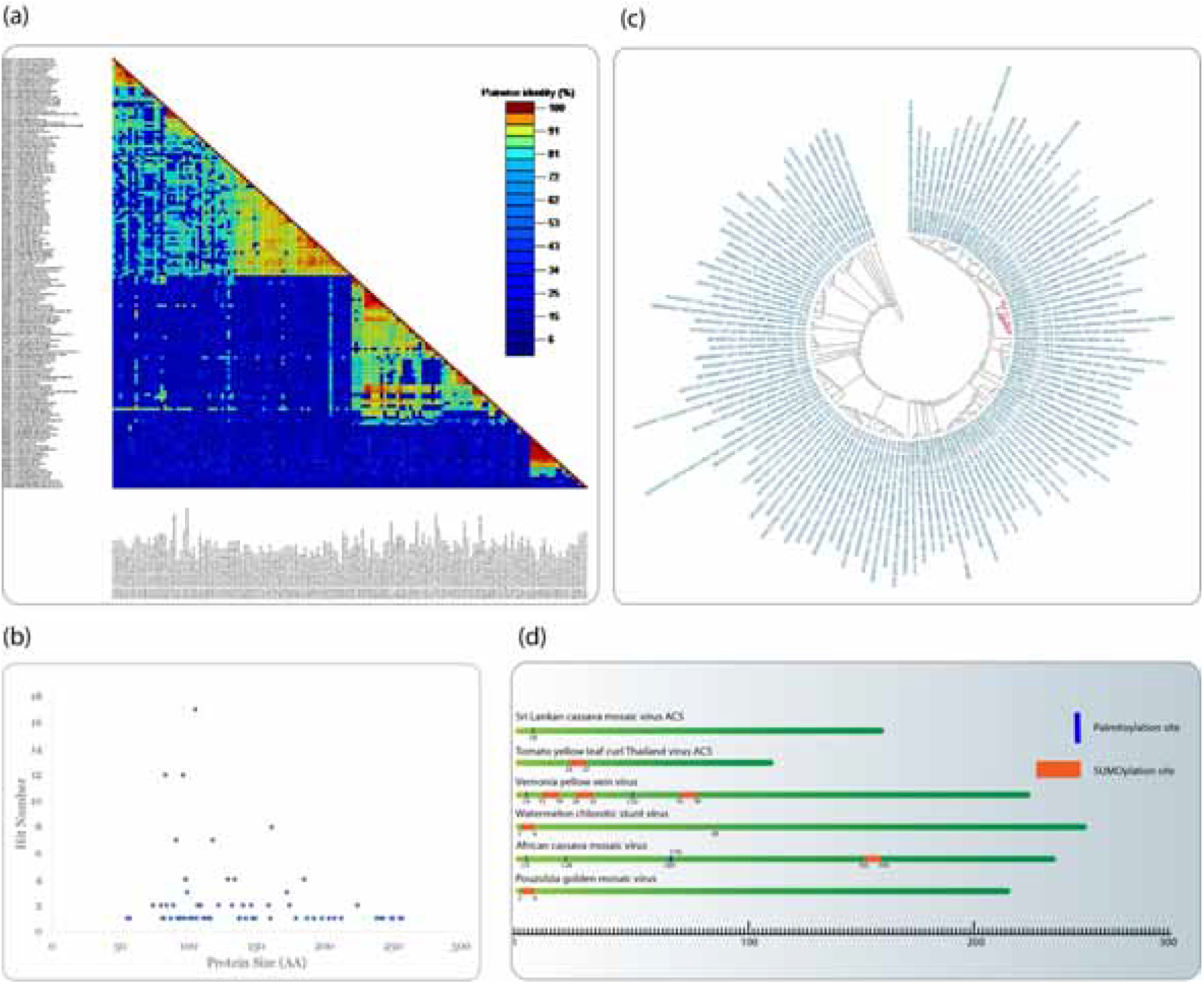
*In-silico* analysis of AC5 proteins. (a) Pairwise sequence identity comparison of begomoviral AC5/C5 protein sequences from database using sequence demarcation tool. Colour code demonstrates the percentage similarity among AC5/C5 sequences. (b) Distribution of differences in amino-acid sequence length between AC5/C5 proteins. (c) A Circular Dendrogram of 130 begomoviral AC5/C5 protein sequences based on UPGMA using MEGA 7. Red clade indicates group with SLCMV AC5 and its closest sequences. (d) Analysis of conservation of functional motifs in AA sequences closely related SLCMV AC5. Left side displays percentage identity with SLCMV AC5 and motif conservation is highlighted in red or blue. Numbers indicate the amino acid position of motif site.

Since these AC5/C5 sequences are closest to SLCMV AC5, we analysed these proteins further, using *in silico* methods, for any predicted post translational modifications which might play an important role during viral infection. The analysis indicated that the AC5 sequences lack myristoylation and ubiquitination sites but contain several palmitoylation and sumoylation sites (Fig 1d).

### AC5 is essential for SLCMV infection

To investigate whether AC5 plays a crucial role in SLCMV infection in *N. benthamiana*, SLCMV infectious clone with a completely mutated AC5 open reading frame (ORF) was utilised. Since the AC5 gene overlaps the coat protein (CP) ORF but is transcribed in the opposite direction, the start codon and all four AUG codons, potentially capable of initiating translation were mutated without affecting the sequence of amino acids of the CP (Fig 2a). The clone containing the mutated version of the AC5 was named AC5m. Furthermore, to make AC5 available ectopically, AC5 was cloned in the PVX based vector (PVX from here on, Chapman, Kavanagh and Baulcombe, 1992) and was validated (Supplementary data Fig S1).

**Figure 2.**
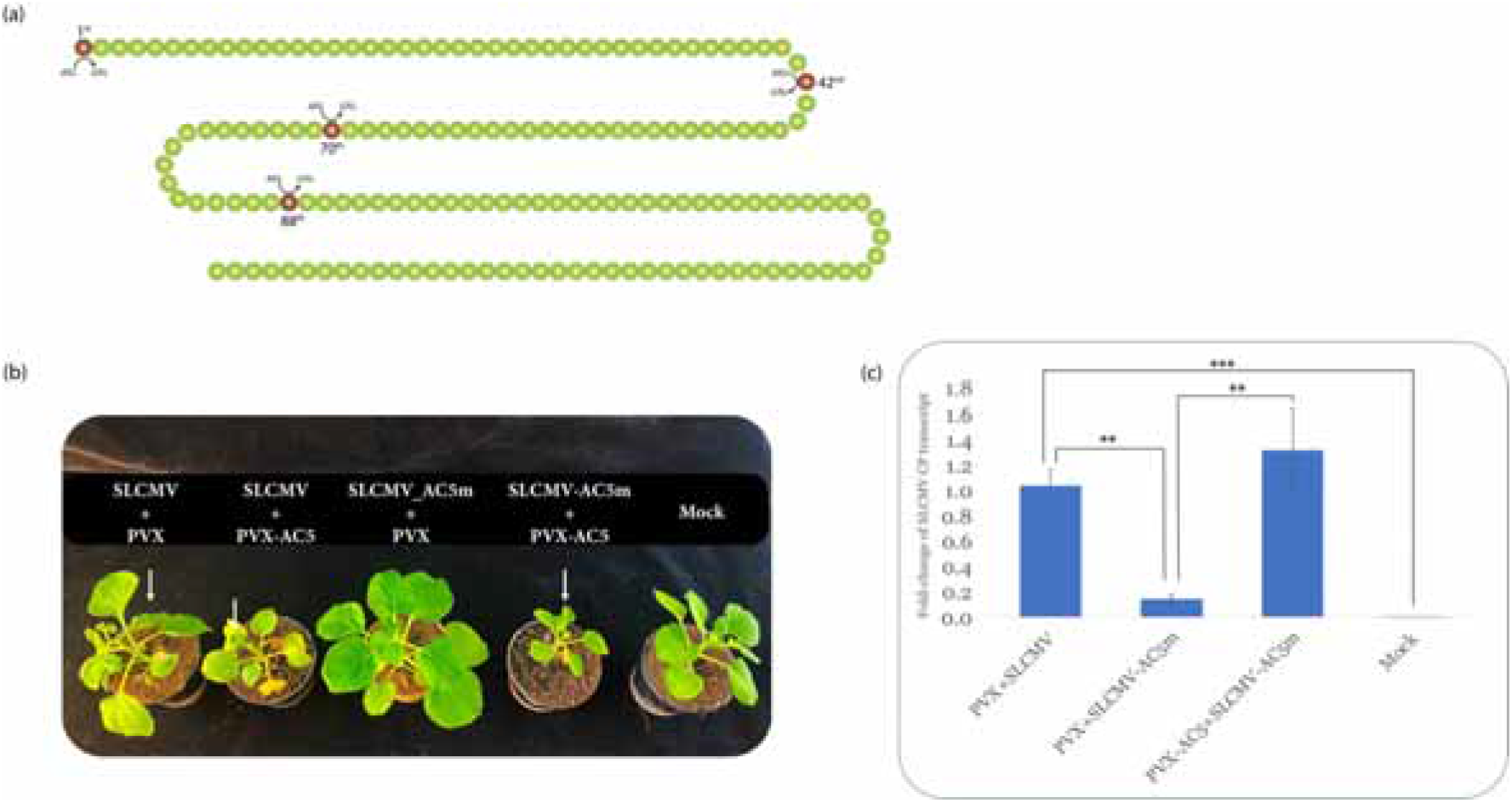
(a) Illustration of mutation sites in AC5 gene. Position of mutation site is respresented. Each sphere represents an amino acid. (b) Co-infection of SLCMV and SLCMV-AC5m with and without PVX-AC5 in *N. benthamiana* plants. Upper panel, Leaf curling symptoms in a *N. benthamiana* plant co-inoculated with SLCMV+PVX, SLCMV+PVX-AC5, SLCMV AC5m+PVX, and SLCMV AC5m+PVX-AC5. The plant inoculated with Agrobacterium cells without viral constructs (Mock) was used as control. The plants were photographed at 15 days post agro-infiltration; white arrow indicates symptomatic leaf (c) Analysis of SLCMV CP expression in the mock- and the co-inoculated plants through qRT-PCR. Error bar represent standard error. P < 0.005 (***), P < 0.01(**), and P < 0.05(*).

SLCMV-AC5m mutant was utilised to analyse infection in *N. benthamiana*. Compared to SLCMV, SLCMV-AC5m was inefficient in inducing symptoms, but when supplied with substantial and continuous AC5 protein through PVX-AC5, it induced significant symptoms (Fig 2b).

For this experiment, 5-week-old *N. benthamiana* was utilised for agroinfiltration. Each combination (SLCMV+PVX, SLCMV+PVX-AC5, SLCMV AC5m+PVX, and SLCMV AC5m+PVX-AC5) was inoculated onto 6 plants. This experiment was carried out for total of three times. As anticipated, symptoms initially developed on plants infiltrated with SLCMV, with or without PVX-AC5 at 12 days post-inoculation (dpi). In the case of SLCMV with a mutant AC5 (SLCMV-AC5m), no symptoms were observed, and the plants resembled those inoculated with empty agrobacterium (mock-inoculated). However, upon supplementation with PVX-AC5 (inoculated with SLCMV + PVX-AC5), at 14 dpi, symptoms became apparent (depicted in Fig 2b, leaf curling highlighted in white arrow), resembling plants inoculated with SLCMV or SLCMV + PVX-AC5. Relative accumulation of SLCMV transcript levels were also quantified in these plants by RT-qPCR using CP primers. Plants infiltrated with SLCMV-AC5m exhibited significantly less SLCMV transcript accumulation, compared to plants infiltrated with SLCMV (Fig 2c). No significant difference was noticed in SLCMV transcript accumulation levels between plants infiltrated with SLCMV and SLCMV-AC5m + PVX-AC5. The emergence of symptoms occurred at variant times, as documented in Table 1 supp data.

Furthermore, viral sequences were purified from symptomatic leaves of plants infiltrated with SLCMV-AC5m + PVX-AC5 and sequenced to validate the maintenance of mutant sites. All mutant sites were found to be intact. To amplify SLCMV AC5 from the replicating SLCMV, specific primers that flank AC5 were utilized, and the resulting PCR products were subjected to sequencing. A total of two plants were infiltrated with SLCMV-AC5m + PVX-AC5, and three symptomatic leaves were selected and sequenced.

### Subcellular localization of AC5 protein

*In-silico* analysis of SLCMV AC5 AA sequence using CSS-Palm (csspalm.biocuckoo.org/online.php) revealed that it contains a putative palmitoylation site on a cysteine residue at 8th position (Fig 1d). S-palmitoylation is a reversible, enzymatic post-translational modification of proteins, characterised by the binding of a palmitoyl chain to a cysteine residue through a thioester linkage. This process of S-palmitoylation has a significant impact on protein behaviour, particularly in terms of its association with membranes, compartmentalisation within membrane domains, trafficking, and constancy. To check whether SLCMV AC5 locates to the cell membrane, AC5 was fused with green fluorescent protein (GFP) in pSITE-2CA (Chakrabarty *et al*., 2007) and was expressed transiently in *N. benthamiana* leaves followed by the analysis of subcellular localization using confocal fluorescence microscopy after 72 hours.

For this experiment, 6-week-old *N. benthamiana* was utilised for agroinfiltration. Leaves were infiltrated with AC5-GFP and Phosphatidylinositol 4,5-bisphosphate-2 tagged with CFP (PIP2-CFP), plasma membrane localization protein and imaged by confocal laser scanning microscopy. Fluorescence from AC5-GFP overlaid fluorescence due to PIP2-CFP, formed abundant bright blue fluorescence along the plasma membrane (PM). The SLCMV AC5 preferentially localized to the PM (Fig 3a). To further confirm the SLCMV AC5 PM localization and exclude the possibility of its apoplastic localization, AC5-GFP was co-expressed with PIP2-CFP and was then subjected to plasmolysis treatment by 20% NaCl before microscopic analysis. Following plasmolysis treatment, the PM was seen to detach from the cell wall (Fig 3b). Green fluorescence from AC5-GFP specifically localized at PM overlaying cyan signal from PIP2-CFP, strongly indicating that of AC5 protein localized to PM. Taken together, the results indicated that SLCMV AC5 was localized in the PM of *N. benthamiana*.

**Figure 3.**
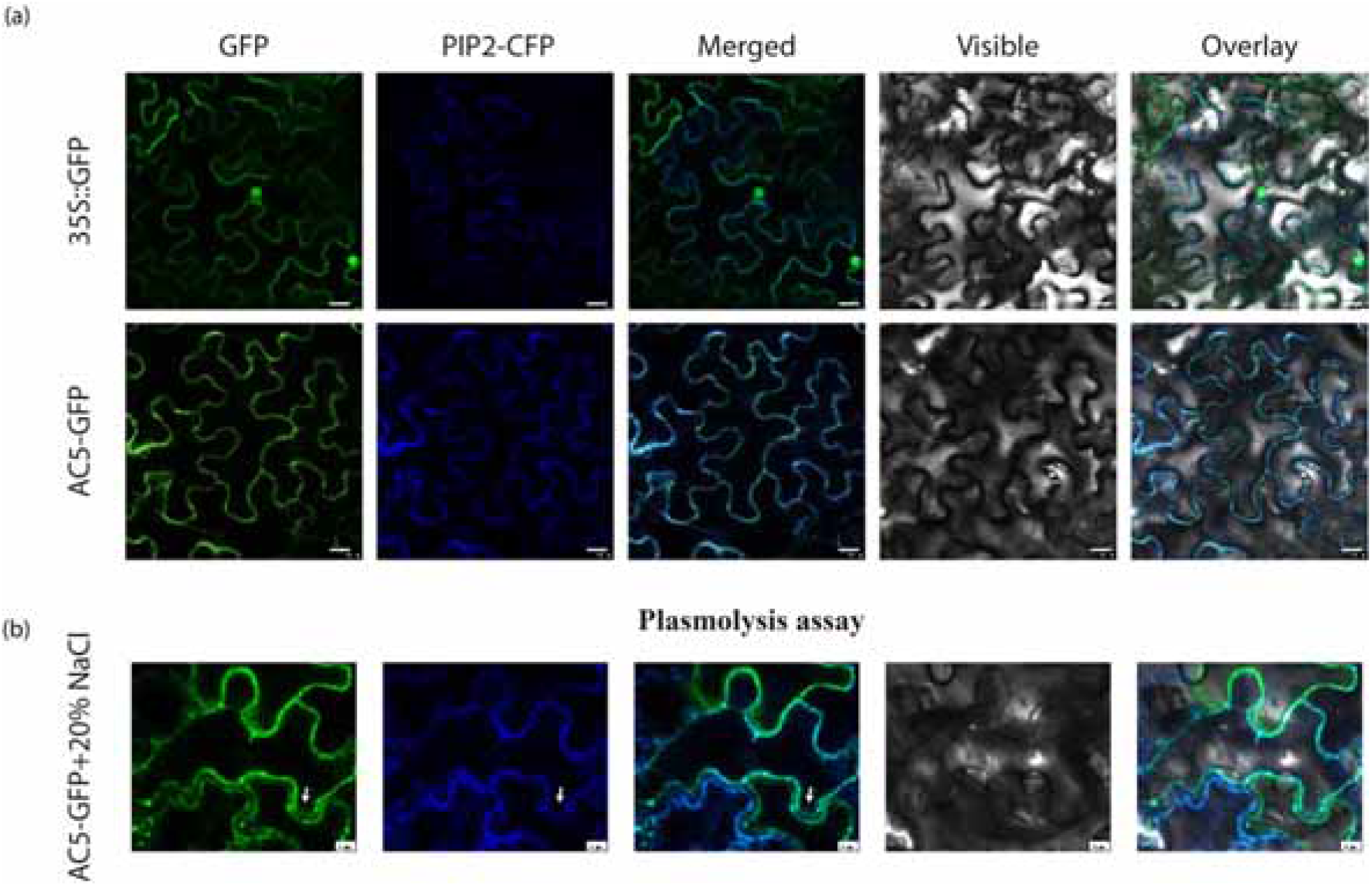
(a) Subcellular localization of AC5 in *N. benthamiana* epidermal cells co-expressed with PM marker PIP2-CFP showing AC5 protein localized in the PM. Scale bars, 10 μM (b) Fluorescence image after plasmolysis showing that confirming that AC5 is membrane protein. The fluorescence of GFP and CFP was monitored at 3 d post-agroinfiltration using confocal laser scanning microscopy.

### SLCMV AC5 interacts with ABI1

To obtain information regarding the potential interactors of SLCMV AC5, yeast two-hybrid screening was performed using an Arabidopsis library encoding biotic stress-induced cDNAs in pGADT7. The initial screening yielded 80 positive colonies that could grow on yeast triple drop-out medium. Out of these, cloned DNA inserts from 20 colonies were subjected to sequencing for their identity and one of them revealed to be encoding for ABA insensitive 1 (ABI1). The interaction between full-length ABI1 and AC5 was confirmed using AC5 as bait (ABI1 in pgGADT7) demonstrating that AC5 interacts with ABI1 in the yeast two-hybrid system (Fig 4a).

**Figure 4.**
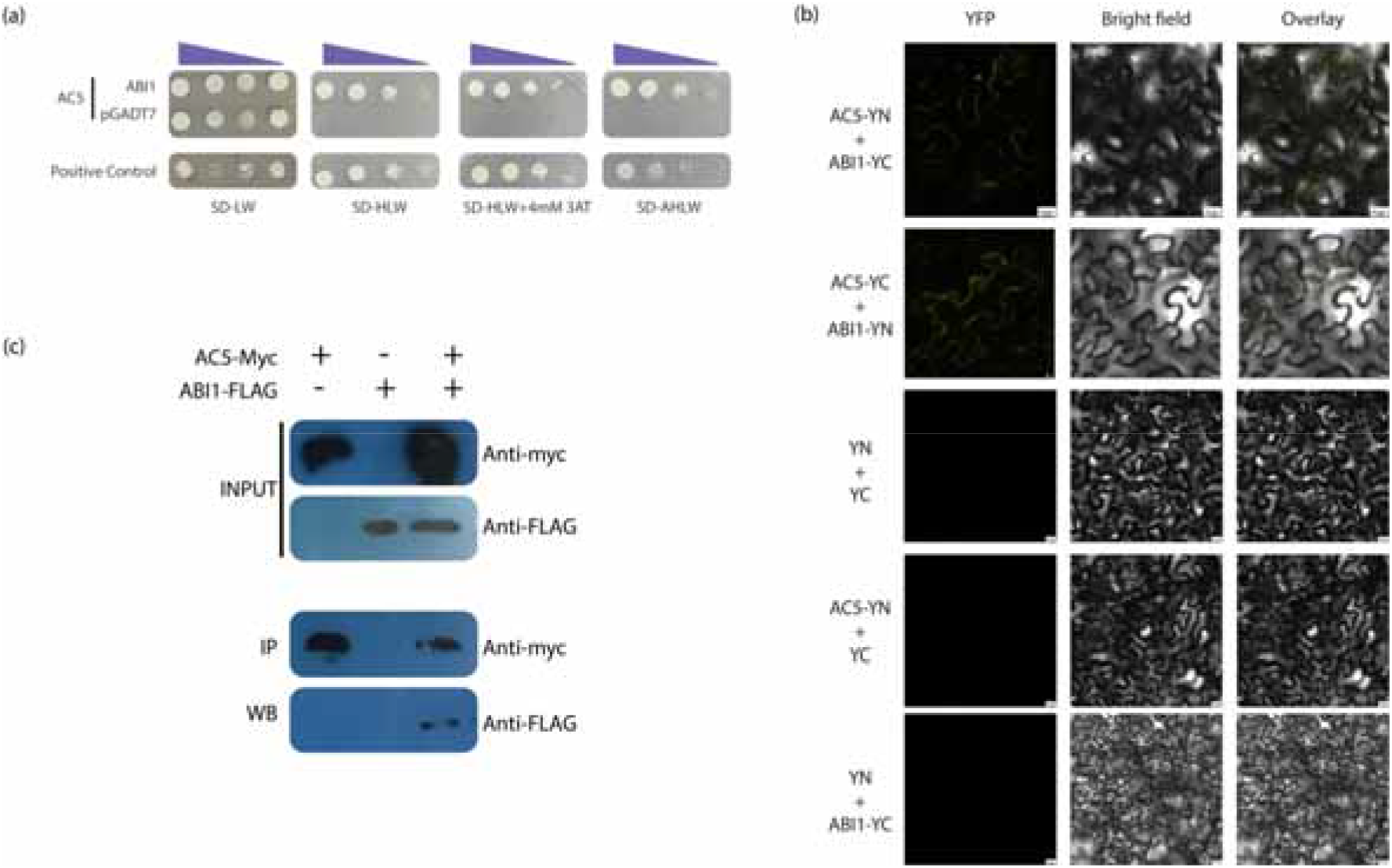
(a) Interaction tests using yeast two-hybrid assays between AC5 and ABI1. Yeast cells with AC5 (BD-AC5) and ABI1 (AD-ABI1) were placed in different liquid concentrations on control medium SD/-Trp/-Leu (SD/LT) and selection medium SD/-Trp/-Leu/-His/-Ade (SD/LTHA). For negative controls, pGADT7 without insert (AD alone), was used. Experiments were performed three times and a representative result is shown. (b) AC5 interacted with ABI1 by BiFC assays in *N. benthamiana* epidermal cells. The recombinant constructs AC5-YN (YFP N-terminal) and ABI1-YC (YFP C-terminal) were agroinfiltrated to *N. benthamiana* leaves. Also, AC5-YC and ABI1-YN was also utilized for validation of interaction. For negative controls, pSPYNE without insert AC5 (YN) and pSPYCE without insert ABI1 (YC) were used. The co-infiltration of YN and YC was also used as a negative control. Fluorescence was recorded 72 hours post infiltration. Experiments were performed 3 times and a representative result is shown. (c) Co-IP assay of AC5 with ABI1. ABI1-Flag and AC5-myc or ABI1-Flag or AC5-myc were transiently expressed in *N. benthamiana* leaves. After 72 h, the total cell lysates were prepared for Co-IP with anti-Myc agarose beads. Then, anti-Myc immunoprecipitates were subjected to Western blot analysis with anti-Flag antibody (lowermost panel). Meanwhile, the total cell lysates were also subjected to Western blot analysis with anti-Flag (middle), and anti-Myc (top panel, for AC5-myc and ABI1-FLAG expression) antibodies. Experiments were performed three times and a representative result is shown.

To verify the reliability of the Y2H results, validation of the interaction in plants was performed using the BiFC assay (Fig 4b). The *ABI1* and *AC5* were fused to the N- and C-termini, respectively, of yellow fluorescent protein gene (*YFP*) to construct fusion vectors (AC5-YN and ABI1-YC), followed by transient infection mediated by agrobacterium of the two fusion proteins in the lower epidermal cells of *N. benthamiana* leaves. At the same time, N-terminus of YFP (YN) was co-expressed with ABI-YC and C-terminus of YFP (YC) was co-expressed with AC5-YN as a negative control. The YFP fluorescence signals of AC5 interacting with ABI1 emerged at 48 h post infiltration along the PM of cell, indicating that their interaction took place at PM (Fig 4b). There was no interaction signal observed for the control combinations of ABI-YC with unfused YN, AC5-YN with unfused YC and empty vectors YC and YN.

Co-immunoprecipitation (Co-IP) experiments were conducted to authenticate the interaction between AC5 and ABI1 in 6-week-old *N. benthamiana* leaves. Three distinct protein combinations, AC5-myc together with ABI1-FLAG, AC5-myc or ABI1-FLAG were infiltrated into *N. benthamiana* leaves through agroinfiltration. Protein extracts of leaves were pulled down with anti-myc antibody. The presence of ABI1 in the same immunoprecipitate was ascertained through the utilization of an anti-FLAG antibody (Fig 4c). These observations validate the interaction between AC5 and ABI1, implying that AC5 could be implicated in the control of ABI1.

### AC5 negatively regulates ABA signalling

To test the effect of AC5 and ABI1 interaction on ABA signalling, three independent AC5 expression lines of Arabidopsis (AC5ox) tagged with myc at C-terminal and driven by CaMV35S promoter were generated. These lines namely L1, L2 and L3 were validated to express AC5 by Western blot (Fig 5a).

**Figure 5.**
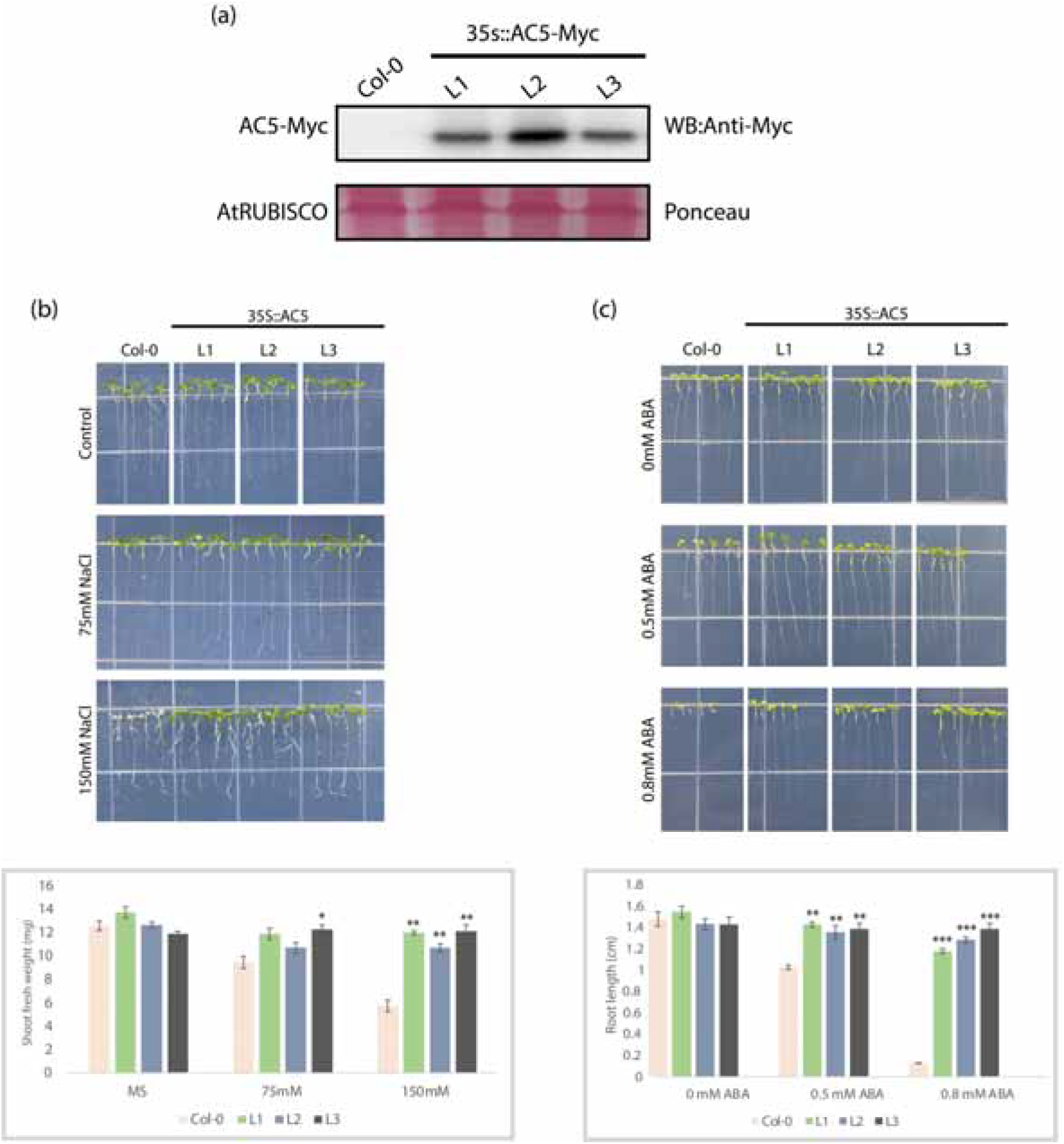
(a) Western blot analysis for AC5-myc. A representative blot of three independent experiments is shown. ABA related phenotypic analysis of 35S::AC5 transgenic lines. (b) Upper panel, phenotypes of 35S::AC5 lines and Col-0 under control (MS) or NaCl treatment (75mM and 150mM), images taken after 6 days. Lower panel, fresh weight of 35S::AC5 transgenic plants and Col-0 Arabidopsis under NaCl treatment. (c) Upper panel, phenotypes of 35S::AC5 lines and Col-0 under control (MS) or ABA treatment (0.5mM and 0.8mM), images taken 6 days of the transfer of 5-day-old seedlings. Lower panel, the root lengths of 35S::AC5 transgenic plants and Col-0 Arabidopsis under ABA treatment. Error bar represent standard error. P < 0.005 (***), P < 0.01(**), and P < 0.05(*).

To evaluate the effect of ABI1-AC5 interaction, certain ABA-related traits were examined in *Arabidopsis thaliana.* ABA governs the expression of numerous salt-responsive genes through transcription factors, whose levels increase in response to salt. Correspondingly, ABI1-mutant Arabidopsis plants manifest heightened sensitivity to salt and osmotic stress during germination, elucidating favorable function of ABI1 in salt tolerance (Suzuki *et al*., 2016). Additionally, ABA suppresses germination. Accordingly, the growth of AC5ox seedlings was assessed in the presence of salt and ABA.

For salt tolerance assay, Arabidopsis seedlings were grown in ½ MS for 7 d, and then transferred to the hydroponic culture plates with 75mM and 150 mM NaCl. The results show that the AC5ox lines had a better growth advantage compared with the Col-0 plants in the presence of 150mM NaCl (Fig 5b). However, no significant difference was observed in plants grown in 75mM NaCl. Difference in growth between AC5ox lines and Col-0 was reflected in shoot fresh weight, demonstrating that AC5ox lines showed enhanced salt tolerance compared to Col-0.

To further prove that AC5 hinders ABA signalling, AC5ox seedling germination rate was tested in the presence of ABA. ABA is known to cause dormancy in seedlings. AC5ox and Col-0 seedlings were germinated vertically in 0.5mM and 0.8mM ABA and root growth was monitored. Under normal growth conditions in ½ MS, no significant difference in the germination rate and primary root growth was observed between Col-0 and the AC5ox lines. However, with ABA treatment, AC5ox lines displayed a significantly higher proportion of seed germination rate and seedling growth than Col-0 and the primary root growth of AC5ox plants was also hyposensitive to ABA treatment (Fig 5c). Highly significant difference was observed in Arabidopsis root growth germinated in 0.8mM ABA. Overall, these data indicate that AC5 reverses the ABA-induced seed dormancy phenotype.

### AC5 expression suppresses AGO gene expression

Recent studies have demonstrated that ABA regulates the expression of several members of the ARGONAUTE (*ago)* family, and this regulation partially contributes to ABA-mediated resistance against BaMV (Alazem and Lin, 2017). Thus, to confirm if overexpression of SLCMV AC5 in Arabidopsis leads to impairment of ABA signalling and dysregulation of *ago* expression, we measured their expression in AC5ox lines. Interestingly, *ago1* and *ago3* were downregulated in AC5ox lines as compared to Col-0 lines (Fig 6a) but not *ago2* and *ago4*. Collectively, ABA has been reported to positively regulate AGO1 and AGO3 (Alazem and Lin, 2017) and upon AC5-ABI1 interaction, the above regulation may be compromised, due to suppression of ABA signalling.

**Figure 6.**
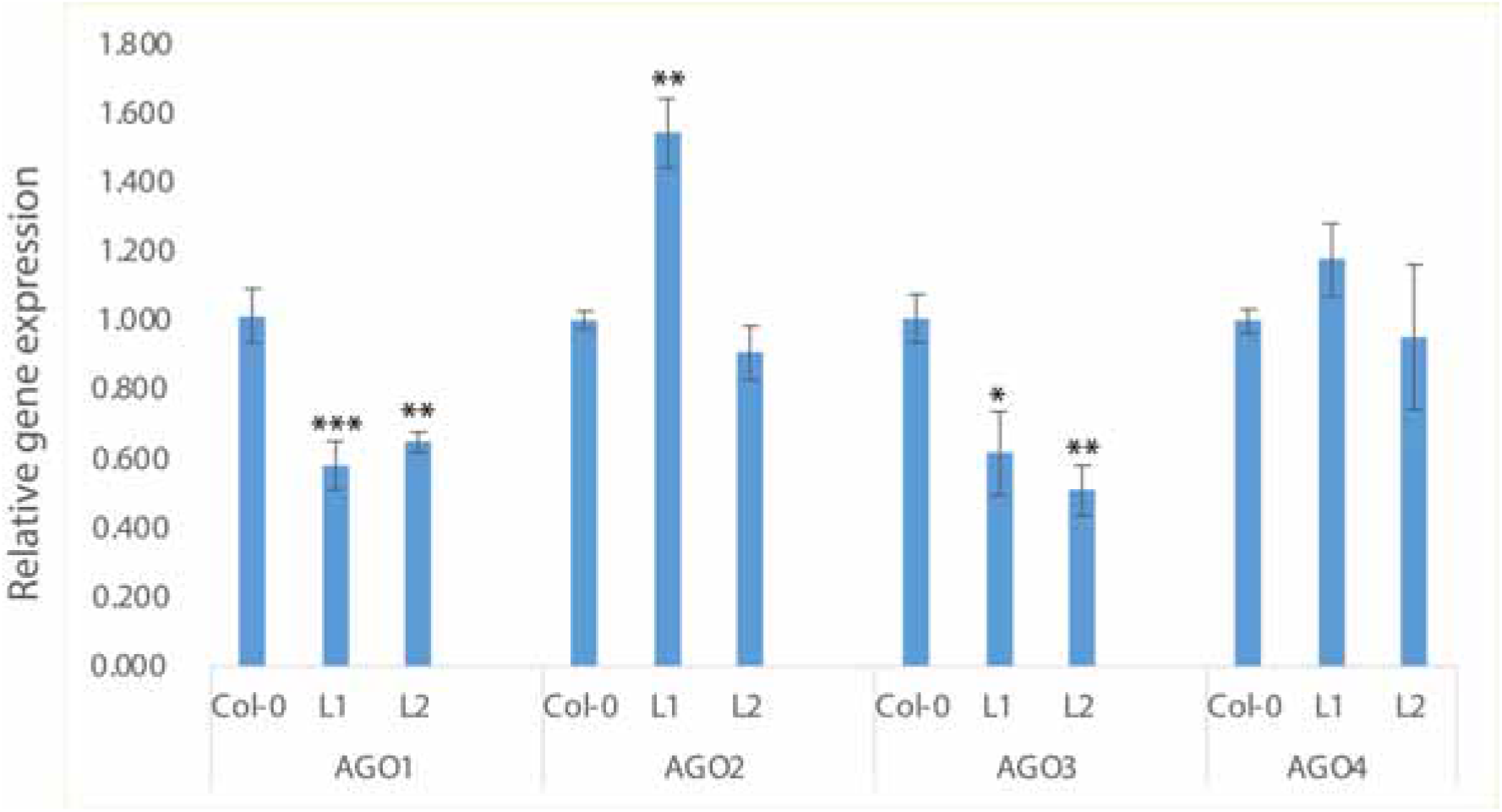
Real-time RT-PCR analyses of the expression of Arabidopsis genes in AC5ox lines and Col-0. RNA was extracted from the leaves of 4-week-old Col-0 and transgenic Arabidopsis plants. The relative amount of the gene transcript was normalized against the expression level of the UBQ gene. AGO1 (AT1G48410); AGO2 (AT1G31280); AGO3 (AT1G31290); AGO4 (AT2G27040). Error bar represent standard error. P < 0.005 (***), P < 0.01(**), and P < 0.05(*).

### Silencing of PP2C decreases SLCMV accumulation in *N. benthamiana*

In Arabidopsis, the *ABI1* encodes a member of the 2C class of protein serine/threonine phosphatases (PP2C). There are multiple orthologs of this gene in *N. benthamiana*. Upon checking the genome sequence, six orthologs of *ABI1* were found in *N. benthamiana* with sequence ID Niben101Scf00769g00012, Niben101Scf03584g03004, Niben101Scf04040g06005, Niben101Scf00611g08002, Niben101Scf09130g05009 and Niben101Scf09170g00002.

To assess how the above genes might contribute to SLCMV infection, a 200 bp long conserved region was selected and cloned in tobacco rattle virus (TRV) based virus-induced gene silencing (VIGS) vector to generate TRV::Anti-PP2C. NbPP2C was silenced in *N. benthamiana* followed by inoculation with SLCMV. Such plants were monitored for symptom development and viral DNA accumulation.

TRV with phytoene desaturase gene (TRV::PDS), obtained from Prof. Arun Kumar Sharma at DPMB, UDSC (Department of Plant Molecular Biology, University of Delhi South Campus), was utilized as the positive control, with the PDS cloned from tomato. The silencing of *pds* produces a typical white color in the emerging leaves that is the result of photo-bleaching, which occurs in the absence of the gene product.

*N. benthamiana* plants were co-inoculated with SLCMV + TRV::Anti-PP2C, negative control SLCMV + TRV and positive control TRV::PDS. Whitening of leaf lamina in newly emerged leaves was observed in PDS-silenced plants at 15 dpi, suggesting a suppression of PDS gene expression (Fig 7a). At 15 dpi, real-time RT-PCR showed that silencing of NbPP2C mRNA levels was to the extent of 60% (Fig 7b).

**Figure 7.**
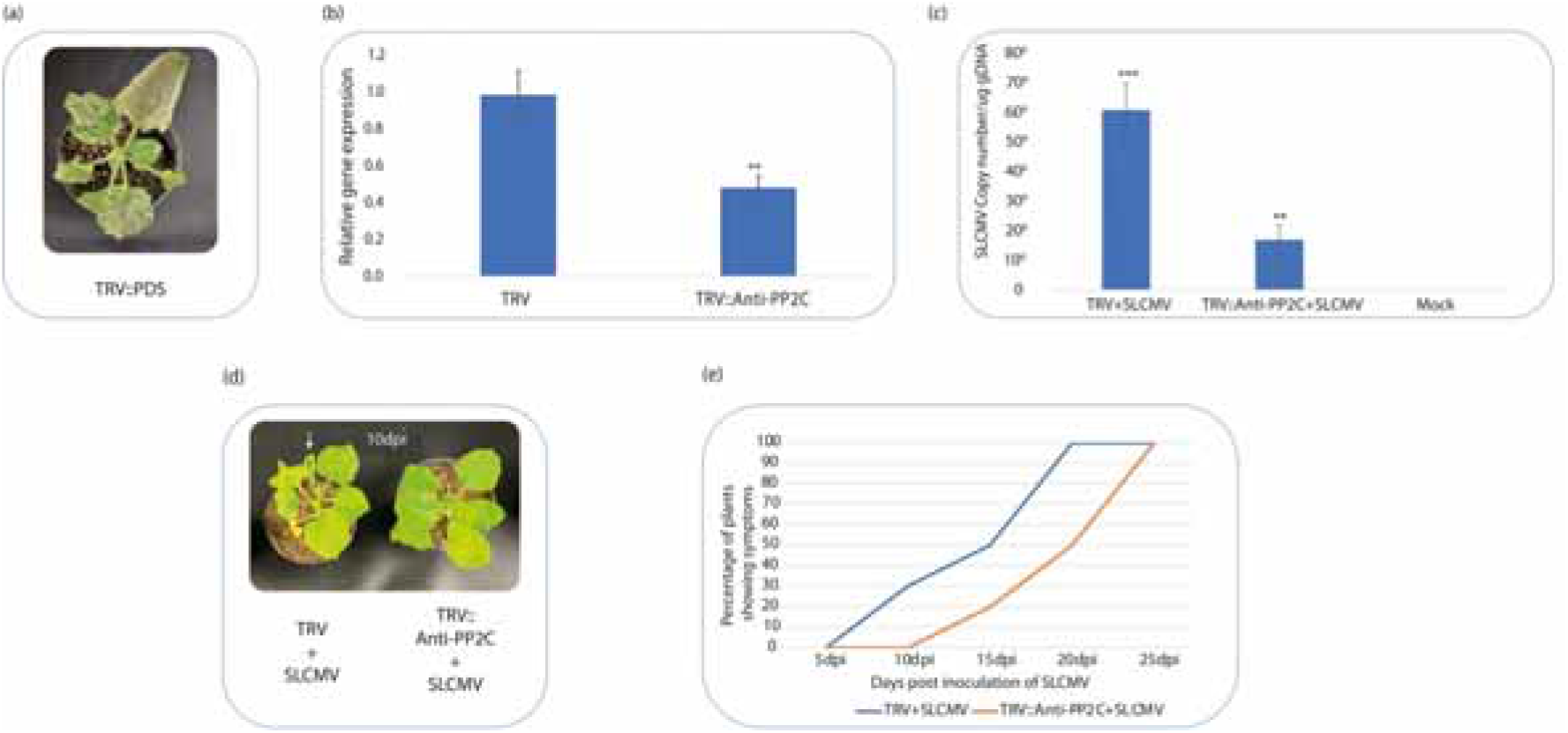
Virus-induced gene silencing in (a) plants infected with TRV vectors of PDS (positive control) (b) RT-qPCR analysis of relative expression of PP2C gene in systemic leaves harvested from the TRV+SLCMV-inoculated plants or the TRV::Anti-PP2C+SLCMV-inoculated plants after 15 days. PP2C transcript levels in TRV+SLCMV-inoculated plants were used as controls, actin was used as the reference gene. Error bars indicate the standard error. The averages presented are based on three independent replicates. (c) SLCMV DNA copy numbers in the systemic leaves harvested from the TRV+SLCMV-inoculated plants or the TRV::Anti-PP2C+SLCMV-inoculated plants after 15 days. (d) plants co-infected with SLCMV/TRV (left, negative control) and SLCMV/TRV::Anti-PP2C (right) in *N. benthamiana*. Symptom of curled leaf after 10dpi is highlighted by white arrow (e) Graph representing comparison of symptom development in plants infected with TRV+SLCMV and TRV::Anti-PP2C+SLCMV (n=20) Error bar represent standard error. P < 0.005 (***), P < 0.01(**), and P < 0.05(*).

Both, plants inoculated with infectious SLCMV construct, followed by infiltration with TRV2 vector (negative control) and TRV::Anti-PP2C showed a slight leaf curl symptom but in case of the latter, symptoms appeared at 15 dpi whereas former displayed symptoms at 10 dpi (Fig 7d and 7e). However, real time PCR revealed SLCMV DNA accumulation to be 3-fold less in PP2C-silenced plants as compared to non-silenced plants (fig 7c). In other words, SLCMV DNA accumulation was significantly lower in NbPP2C-silenced plants compared to control plants. This was also observed in delaying symptom development in NbPP2C silenced lines (Fig 7e).

## Discussion

A multitude of geminiviruses, including MYMIV, tomato leaf deformation virus, ToCMoV, and WmCSV, contain the C5/AC5 ORF in the homologous strand of their genomes (Kheyr-Pour *et al*., 2000; Fontenelle *et al*., 2007; Li *et al*., 2015, 2021). Upon downloading and comparing the sequences of 150 randomly selected AC5/C5 proteins, the results showed that it is highly variable in terms of length as well as sequence identity. This strongly suggests that AC5 may encode a protein with varying functions in different begomoviruses. For example, although the MYMIV AC5 has been shown to be a virulence factor with a specific role in pathogenicity (Li et al., 2015), for WmCSV and ToCMoV, AC5 is not required for infection (Kheyr-Pour *et al*., 2000; Fontenelle *et al*., 2007).

To validate if AC5 is required for SLCMV, mutational analysis demonstrated that the SLCMV defective in AC5 expression (SLCMV-mAC5) failed to cause clear disease symptoms and accumulated much less viral DNA. Ectopic expression of AC5 using PVX allowed SLCMV-mAC5 to accumulate in plants and cause symptoms. Consequently, we conclude that SLCMV AC5 protein is important for symptom development and virus accumulation.

Subsequently, a phylogenetic analysis was conducted to evaluate the closely related proteins, elucidating AC5 proteins from African cassava mosaic virus, vernonia yellow vein virus, TYLCTHV, and watermelon chlorotic stunt virus. Accordingly, a study of the conserved functional post-translational-modification (PTM) motifs among these sequences revealed the presence of a palmitoylation site in a subset of them. Palmitoylation is a reversible PTM, wherein a C16-carbon saturated fatty acyl chain is covalently joined to the cytoplasmic cysteine residues; this phenomenon is catalyzed by palmitoyl-acyl transferases and reversed by protein palmitoyl thioesterases (Hemsley and Grierson, 2008), the mechanisms of which remain obscure. Palmitoylation of cytoplasmic proteins has been extensively demonstrated to modulate the interaction of these soluble proteins with specific cellular membranes or membrane regions (Hemsley and Grierson, 2008). We speculate conservation of palmitoylation site in closely related sequences might be related to functional conservation.

In this study, we have found that SLCMV AC5 protein is localized in the PM. Additionally, to counterbalance the supposition that it could be an apoplastic protein, a plasmolysis study was conducted. These investigations yielded evidence that SLCMV is likely to be a PM localized protein unlike other C5/AC5 which localize in cytosol and nucleus for instance ALCScV C5 and squash leaf curl China virus AC5 (Li *et al*., 2021; Wu *et al*., 2022). These differing localizations may reflect variations in their functions.

ABA is well-known for its role in plant responses to abiotic stress and for its multifaceted roles in plant–pathogen interactions. We demonstrate that SLCMV AC5 physically interacts with the central negative regulator of ABA signalling, ABI1, probably resulting in the perturbation of the ABA-mediated defence signalling pathways. To further confirm if this interaction is significant, we assessed phenotypes associated with ABA in AC5-expressing transgenic Arabidopsis plants. These lines were validated for expression of AC5 by western blotting. However, to verify if it is a pathogenicity element and does influence defence associated genes, the transcriptome of defence related genes was analyzed and as expected, there was significant differential expression of defence related genes (Supplement data Fig S2). These AC5-expressing lines were checked for the alteration of functions associated with ABA such as salinity tolerance assay and a germination assay. Results demonstrated that the AC5-expressing plants were tolerant to 150 mM NaCl salt and germinated in the presence of 0.8 µM ABA, sufficient to inhibit germination, suggesting that AC5 renders the plants ABA-insensitive. This interaction may imply a potential method of suppressing ABA-dependent defence pathways to facilitate the propagation of SLCMV.

ABA, a phytohormone of fundamental significance, mediates a plethora of plant growth and development processes, encompassing seed germination and fruit maturation, as well as varied abiotic stresses (e.g., drought, salinity, and hydric stress; Finkelstein, Gampala & Rock, 2002; Tuteja, 2007). Moreover, earlier studies have elucidated the essential role of ABA in the context of biotic stresses (Ton, Flors and Mauch-Mani, 2009). The induction of resistance to the fungal pathogen *Cochliobolus miyabeanus* (de Vleesschauwer *et al*., 2010) and the promotion of susceptibility to *Magnaporthe oryzae* in the same host plant by ABA have been documented (Xu *et al*., 2013). The role of ABA in plant immunity varies depending upon the stage of infection. In the initial stages, ABA can act in a positive manner, fostering defence via stomatal closure and callose deposition, but its influence is deleterious when the infection reaches a later stage, as it impedes signalling from SA and JA (Ton, Flors and Mauch-Mani, 2009). Studies into the impact of ABA on viral infections have been much less extensive than those involving bacteria and fungi, yet the existing evidence shows that ABA may enhance resistance. For instance, the effect of TMV on ABA accumulation has been observed (Whenham *et al*., 1986) with ABA-enhanced callose deposition seen to limit virus movement (Rezzonico *et al*., 1998). More recently, Arabidopsis–BaMV interactions demonstrated that ABA induces plant resistance by regulating AGO2 and AGO3 (Alazem *et al*., 2017). Similarly in our study, quantification of *ago* expression in AC5-expressing Arabidopsis revealed a marked decrease in expression of *ago1* and *ago3* genes (fig 6a). Nonetheless, no considerable variation was detected in *ago2* and *ago4*. This might be attributed to inhibition of ABA signalling by AC5-ABI1 interaction. Decrease in *ago* expression might lead to drop in RNAi against virus leading to increased virulence.

However, the mechanism by which the AC5-ABI1 interaction results in the suppression of ABA signalling remains enigmatic. There are various hypotheses that could account for this phenomenon, such as the possibility that AC5 prevents ABA-mediated phosphorylation inhibition of ABI1 or that this interaction causes conformational changes in ABI1 that result in the perpetual sequestration of SnRKs by ABI1. Further investigation is necessary to fully comprehend this process. This study presents the first evidence of a geminiviral AC5 interacting with ABI1 and its role in decreasing ABA signalling.

To further elucidate the part of PP2Cs in geminiviral pathogenesis, we employed VIGS-mediated gene silencing to suppress PP2C expression and then study symptomology and viral titer development in SLCMV-infected *N. benthamiana* plants. Suppression of PP2C expression by 60% resulted in a 3-fold reduction of SLCMV titers by15 dpi. This may be due to augmented ABA signalling and its defence related roles.

In summary, our results demonstrate the essentiality of the SLCMV AC5 protein in promoting infection and its interaction with ABI1. This interaction culminates in the disruption of ABA signalling, consequently inducing an ABA-insensitive phenotype in Arabidopsis (Fig 8). This impairment subsequently hampers ABA-mediated defence mechanisms, including the arrest of ABA-related defence pathways. Notably, the induction of AGO genes, a pivotal mechanism through which the ABA defence pathway operates, was found to be marginally compromised in Arabidopsis lines expressing AC5. This strategic interference by AC5 in ABA-mediated defence pathways exemplifies a counter-defensive approach that establishes a favourable environment for viral proliferation.

**Figure 8:**
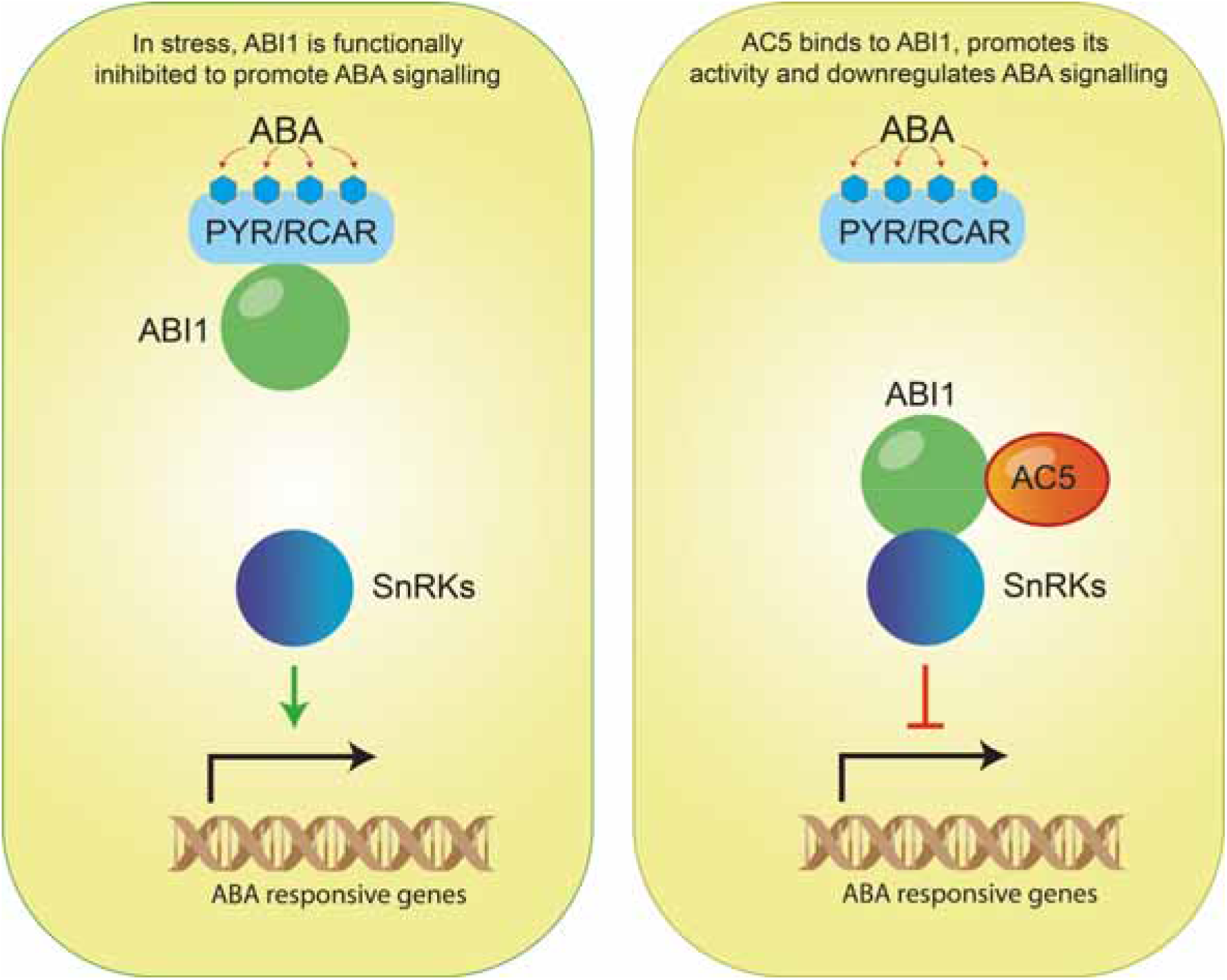
Schematic representation of AC5 mediated inhibition of ABA signalling. AC5 exerts a negative impact on ABA signalling by interacting with ABI1. When subjected to stress, in the presence of ABA, PYR1 forms a complex with ABA, subsequently binding to and restraining ABI1. Consequently, this interaction liberates SnRKs from their sequestered state alongside ABI1 (Left panel). However, the presence of SLCMV AC5 alters this dynamic by maintaining ABI1’s inhibition on ABA signalling, thereby inducing desensitization of the signalling pathway to ABA (Right panel).

## Acknowledgement

ID would like to acknowledge the J. C. Bose Fellowship (SB/S2/JCB-057/2016) from the Science and Engineering Research Board, Government of India, whose funds were utilized for conducting this study. RK would like to acknowledge Research Fellowship by the Department of Biotechnology, Government of India (DBT/JRF/BET-17/I/2017/AL/455) during the tenure of this work. Funds made available to ID under the Faculty Research Grant, University of Delhi are also acknowledged. Authors acknowledge the facilities made available under the FIST infrastructure Grant to the Department, which were utilized in this study. The authors would like to express gratitude to Prof Girdhar Kumar Pandey for graciously providing the Arabidopsis yeast two-hybrid library and Prof Arun Kumar Sharma for the TRV vector.

## Author contributions

RK and ID conceived of the project and designed the research; RK performed the experiments and *in silico* work; RK and ID analyzed the data; RK and ID wrote the manuscript; ID acquired the funds; RK and ID edited the manuscript.

